# ENERGY OPTIMIZATION DURING WALKING CAN BE A PRIMARILY IMPLICIT PROCESS

**DOI:** 10.1101/2020.08.21.261057

**Authors:** Megan J. McAllister, Rachel L. Blair, J. Maxwell Donelan, Jessica C. Selinger

## Abstract

Gait adaptations, in response to novel environments, devices or changes to the body, can be driven by the continuous optimization of energy expenditure. However, whether energy optimization is primarily an implicit process—occurring automatically and with minimal cognitive attention—or an explicit process—occurring as a result of a conscious, attention-demanding, strategy—remains unclear. Here, we use a dual-task paradigm to test whether energy optimization during walking is primarily an implicit or explicit process. To create our primary energy optimization task, we used lower-limb exoskeletons to shift people’s energetically optimal step frequency to frequencies lower than normally preferred. Our secondary task, designed to draw explicit attention from the optimization task, was an auditory tone discrimination task. We found that adding this secondary task did not disrupt energy optimization during walking; participants in our dual-task experiment adapted their step frequency toward the optima by an amount similar to participants in our previous single-task experiment. We also found that performance on the tone discrimination task did not worsen when participants were optimizing for energetic cost; accuracy scores and reaction times remained unchanged when the exoskeleton altered the energy optimal gaits. Survey responses suggest that dual-task participants were largely unaware of the changes they made to their gait to optimize energy, whereas single-task participants were more aware of their gait changes yet did not leverage this explicit awareness to improve gait optimization. Collectively, our results suggest that energy optimization is primarily an implicit process, allowing attentional resources to be directed toward other cognitive and motor objectives during walking.

**Summary statement:** People can adapt to energy optimal walking patterns without being consciously aware they are doing so. This allows people to discover energetically efficient gaits while preserving attentional resources for other tasks.

## Introduction

Humans have a remarkable ability to adapt their gait to changing terrains, tasks, and even constraints on their body. When we encounter a steep hill, navigate a crowded space, or carry a heavy load, we change how we walk. Although we often do so with relative ease, the underlying control mechanism is necessarily complex. To coordinate the movements of our limbs, we adjust the time-varying activation of tens of thousands of motor units across hundreds of muscles. In turn, by altering these coordination patterns we choose between different gaits, such as walking or running, and adapt countless gait parameters, such as speed, step frequency, and limb symmetry. Our research group, and others, have recently demonstrated that gait adaptations can be driven by continuous optimization of energy expenditure—when searching the expanse of possible gaits, we often prefer and converge on those that minimize the calories we burn in a given context (Abram et al. 2019; Finley et al. 2013; Roemmich et al. 2019; Selinger et al. 2015, 2019). However, whether energy optimization is primarily an *implicit process*—occurring automatically and with minimal cognitive attention—or rather an *explicit process*—occurring as a result of a conscious, attention-demanding, strategy—remains unclear (Frensch 1998; Kahneman and Egan 2011; Mazzoni and Krakauer 2006). For example, when we encounter a hill, we might implicitly slow our speed and reduce our step rate, without even realizing it (Kawamura et al. 1991; Sun et al. 1996). Or, we might see the steep terrain, judge it looks tiring, and explicitly decide on a strategy to slow down and alter our angle of approach to reduce steepness. Both implicit and explicit processes may be used to reduce energy expenditure, either in isolation or in unison.

Dual-task paradigms have been used to assess to what extent a task is implicit or explicit in nature. Typically, a *primary task* of interest is simultaneously performed with a *secondary task* known to require explicit processing, such as counting backwards or stating the color of text incongruent with the word it spells (Beauchet et al. 2005; Bench et al. 1993; Kahneman 1973; Stroop 1935). The theory underlying this design is that our cognitive attention is a *limited capacity resource*—we can only think and explicitly strategize about so many things at a time (Magill 2011; Schmidt and Lee 2011; Woollacott and Shumway-Cook 2002). Therefore, if the secondary task is sufficiently challenging and the primary task is explicit in nature, performance on one or both tasks will be hindered. Alternatively, if the primary task is implicit in nature, performance decrements should not occur. For example, dual-task paradigms have been used to interrogate the role of explicit control in walking. In able-bodied adults, during unperturbed walking in a predictable environment, walking is primarily an implicit process (Lajoie et al. 1993; Malone and Bastian 2010; Paul et al. 2005; Regnaux et al. 2005). Regardless of the nature of the secondary explicit task, be it counting backward, verbally repeating sentences, or buttoning a shirt, walking performance characteristics, such as speed, step length, and the variability of each, are largely unchanged (Beauchet et al. 2003; Ebersbach et al. 1995; Lajoie et al. 1999; Paul et al. 2005). This is not however the case in all contexts and for all populations.

Dual-task paradigms have been used to demonstrate the enhanced role of explicit control when navigating obstacles during walking or when stepping to defined visual targets like one might encounter on a stone path (Mazaheri et al. 2014; Peper et al. 2012; Sparrow et al. 2002; Weerdesteyn et al. 2003). They have also been used to demonstrate that in children, older adults, and individuals with cognitive impairments, even unperturbed straight-line walking can involve significant explicit control, evidenced by slowing gait speeds and increased variability under the demands of a secondary task (Beauchet et al. 2003; Hagmann-von Arx et al. 2016; Lajoie et al. 1999; Li et al. 2000; Montero-Odasso et al. 2012; Theill et al. 2011). Dual-task paradigms are a tool to probe the nature of explicit control during movement and have been used extensively in walking contexts.

While dual-task paradigms have been used for decades to probe the nature of various well-learned motor tasks like walking, they have only recently been applied to the *adaptation* of motor tasks (Conradsson et al. 2019; Malone and Bastian 2010; Taylor et al. 2014; Taylor and Thoroughman 2007). Motor adaptation, where a well-learned movement is modified in response to a new context through trial and error, has long been assumed to be an implicit process (Benson et al. 2011; Masters et al. 2008; Mazzoni and Krakauer 2006; Willingham 1998). For example, in canonical force-field paradigms, where forces from a robotic manipulandum alter limb dynamics during reaching, a common understanding is that adaptation is driven by sensory-prediction errors that update an internal model (or stored prediction) of the task dynamics. (Shadmehr et al. 2010; Shadmehr and Mussa-Ivaldi 1994). This recalibration was thought to be primarily automatic, occurring below the level of conscious control. However, recent work has revealed that explicit processes can play a significant role in adaptation (Conradsson et al. 2019; Malone and Bastian 2010; Taylor et al. 2014; Taylor and Thoroughman 2007). In one experiment, Taylor and Thoroughman (2007) had participants perform a tone discrimination task (secondary explicit task) while adapting to perturbations from a novel force-field during reaching (primary task). They found participants’ ability to correct arm position during a given movement was not affected, but adaptation from one reach to the next was (Taylor and Thoroughman 2007). This implies that within-movement feedback control may be primarily implicit, but that movement-to-movement error corrections and the updating of predictive control involves explicit strategy (Taylor and Thoroughman 2007). In later visuomotor adaptation experiments, Taylor et al. (2014) confirmed these findings and were able to decouple the contribution and time course of implicit and explicit processes during adaptation by asking participants to verbalize their aiming direction (state their explicit strategy) at the onset of each reach. Evidence from walking paradigms have provided further evidence that motor adaptation can in fact involve explicit strategy. In split-belt treadmill walking paradigms, where participants adapt to belts travelling at different speeds under each foot, explicit secondary tasks can disrupt adaptation, particularly in older adults (Conradsson et al. 2019; Malone and Bastian 2010). Current understanding is that motor adaptation, whether in discrete upper-arm reaching tasks or continuous lower-limb walking tasks, can involve both implicit and explicit processes.

Here, we use a dual-task paradigm to test whether *energy optimization* during walking is primarily an implicit or explicit process. We define energy optimization, our primary task of interest, as the process of adapting one’s gait to minimize metabolic energy expenditure. To study the energy optimization process, we leverage our previous experimental paradigm where robotic exoskeletons are used to shift people’s energetically optimal step frequency to frequencies lower than normally preferred (Selinger et al. 2015, 2019). We evaluate performance in this task by adaptation toward the optima, measured by decreases in step frequency. We have previously shown that people adapt to energy optimal step frequencies when performing only this task (in a single-task context). Here, we add a secondary tone discrimination task to this primary energy optimization task. This explicit secondary task requires that participants indicate whether a current audio tone is of higher or lower frequency than the previous tone. Performance in this task is evaluated in terms of accuracy (correct responses) and reaction time (time to respond). One hypothesis is that energy optimization during walking is primarily an implicit process and performance in both tasks will be maintained. This would be consistent with the more traditional perspective that the control of well-learned movements, and the motor adaptation of these movements, are largely automatic and occur below the level of conscious control (Lajoie et al. 1993; Malone and Bastian 2010; Paul et al. 2005; Regnaux et al. 2005; Shadmehr et al. 2010). An alternative hypothesis is that energy optimization is primarily an explicit process and performance on one or both tasks will deteriorate. This would be consistent with the more recent findings that motor adaptation, in both reaching and walking paradigms, can result from conscious execution of an explicit strategy (Conradsson et al. 2019; Malone and Bastian 2010; Taylor and Thoroughman 2007).

## Methods

### Participants

We performed testing on a total of 11 healthy adults (7 female, 4 male) with no known gait, cardiopulmonary, or cognitive impairments. Simon Fraser University’s Office of Research Ethics approved the protocol, and participants gave their written, informed consent before testing.

### Primary energy optimization task

To create a task where participants had to adapt their gait in order to minimize energy expenditure, we leveraged our previous paradigm where robotic exoskeletons are used to shift people’s energetically optimal step frequency. We have previously shown, in a single-task context, that people adapt toward energy optimal step frequencies (Selinger et al. 2015, 2019). We used custom software to measure and control the magnitude of the resistive torque applied to the knees in real-time at 200 Hz (Simulink Real-Time Workshop, MathWorks). In our current experiment, all participants experienced a ‘penalize-high’ control function where the resistive torque, and therefore added energetic penalty, was minimal at low step frequencies and increased as step frequency increased (Fig. 1B) (Selinger et al. 2015). This function reshapes the *energy landscape*—in this case the relationship between step frequency and energetic cost—creating a positively sloped energetic gradient at the participants’ naturally preferred step frequency, and an energetic minimum at a lower step frequency (Fig. 1C). To implement this control function, we made the commanded resistive torque to the exoskeleton proportional to the participants’ step frequency measured from the previous step (Fig. 1A). To measure step frequency at each step, we calculated the inverse of the time between foot contact events, identified from the fore–aft translation in ground reaction force centre of pressure from the instrumented treadmill (FIT, Bertec Inc.). We sampled ground reaction forces at 200 Hz (NI DAQ PC1-6071E, National Instruments Corporation). When commanding step frequency to the participants, we used a custom auditory metronome (Simulink Real-Time Workshop, MathWorks). Full details about the exoskeleton hardware, controller and paradigm can be found in our previous papers (Selinger et al. 2015, 2019). To measure participants’ resulting energy expenditure throughout the protocol, we used indirect calorimetry (VMax Encore Metabolic Cart, VIASYS®).

**Figure 1:**
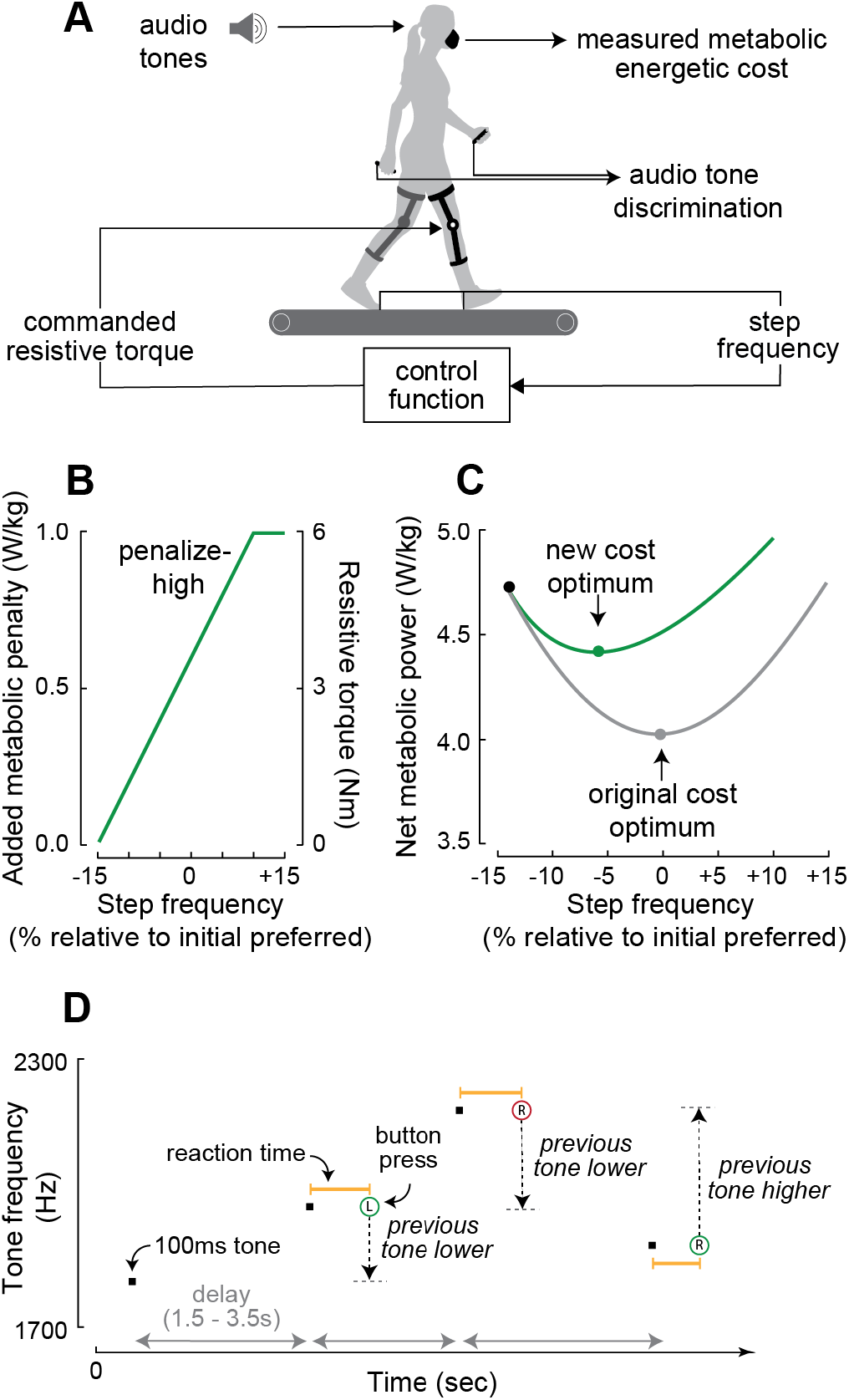
Dual-task experimental design. **A:** To create the primary energy optimization task, a control function commands resistive torques to the knee exoskeletons that are proportional to step frequency, making higher step frequencies energetically costly and lower step frequencies less costly. To create the secondary tone discrimination task, audio tones are presented, and the participant must indicate if the frequency of the current tone is higher or lower than the preceding tone by pressing a button held in the right or left hand, respectively. **B:** Design of the penalize-high control function. **C:** Schematic energetic cost landscape of the penalize-high control function (green) and the original cost landscape (grey). **D:** In the secondary task, we used custom software to output a steady stream of 100 ms audio tones (black squares) with a frequency between 1700 and 2300 Hz. The time between tones randomly varied from 1.50-3.50s (horizontal grey arrows). Left-hand and right-hand button presses are represented by circles encompassing a **L** or **R**, respectively. A button press circle colored green indicates a correct response, while red indicates an incorrect response. The dashed vertical arrows indicate the difference in frequency between the current tone and the preceding tone. Reaction times, from onset of tone to button press, are indicated by the horizontal yellow lines.

### Secondary tone discrimination task

To create a secondary explicit task, we used a *one-back* audio tone discrimination task (Fig 1D). In this task, participants listened to a stream of auditory tones and continually distinguished if the present tone was of higher or lower frequency than the tone immediately preceding it (one-back) (Kane et al. 2007). In pilot testing (n=2), under natural walking conditions (no exoskeleton), we also explored a simpler *paired-tone* task, where participants distinguished the frequency between two tones presented sequentially and can then discard them from memory (Taylor and Thoroughman 2007), as well as a more complex *two-back* task, where the participants must continually distinguish if the present tone is of higher or lower frequency than the second from last tone preceding it (two-back) (Kane et al. 2007). Consistent with findings from Taylor and Thoroughman (2007), we found that the paired-tone task may not be challenging enough to sufficiently tax the explicit cognitive process. Average scores were consistently above 90%. Conversely, we found the two-back task was likely too challenging (correct response rates only slightly higher than 50% chance rate), risking participant disengagement. We settled on the one-back task, for which average responses were just above 80% in piloting.

To implement the one-back tone discrimination task, we output the stream of audio tones to a speaker using custom software (Matlab 2013b, MathWorks) (Fig. 1D). We made the duration of each tone 100ms, while the time between tones ranged from 1.50-3.50s, chosen randomly from a uniform distribution (Taylor and Thoroughman 2007). To output tones of continuously varying frequencies, we created a three-tone loop. The frequency of the first tone in the three-tone loop was randomly selected from a uniform distribution (2000 Hz ± 150 Hz). The second and third tones in the three-tone loop occurred at a frequency ± 150 Hz of the first tone, chosen randomly from a uniform distribution. The participants held a thumb activated push-button in each hand and we gave them the following instructions:

*You will be conducting a one-back audio discrimination task over the duration of each trial. That means you will listen to a stream of tones and compare the tone you just heard to that immediately before it. You are comparing tones in terms of higher or lower sound.*

*Once you have determined that the tone you just heard was higher or lower than that immediately preceding, indicate your response via a button press. A left button press means lower and a right button press means higher. Just remember, left equals lower.*

We collected button press analog signals, as well as tone frequency, timing and duration through a data acquisition board (BNC-2110, National Instruments) using a custom software script (Matlab 2013b, MathWorks). To ensure that participants understood the instructions and could adequately execute the secondary task, they practiced during a one-minute sample of the tone discrimination task, prior to our experimental protocol, while standing.

### Experimental protocol

We replicated the protocol of our previous experiment (Selinger et al. 2019), but with the addition of the secondary tone discrimination task. This was done to allow us to directly compare dual-task and single-task results. The protocol consisted of four testing periods: Baseline Period, Habituation Period, First Experience Period, and Second Experience Period (Fig. 2). Participants performed the secondary tone discrimination task throughout the entirety of all four periods while walking on the treadmill at 1.25 m/s. We provided 5-10-minute rest periods between each period. During the Baseline Period, participants walked for 12 minutes with the exoskeleton controller turned off (Fig. 2A). We used this period to determine participants’ *initial preferred step frequency* under natural conditions, calculated as the average step frequency during the last 150 seconds of the period. During the Habituation Period, to familiarize participants with walking at a range of step frequencies while completing the tone discrimination task, we instructed participants to match their steps to both high and low frequency metronome tempos (+10% and −15% of their initial preferred step frequency, respectively) over the course of 18 minutes (Fig. 2B). The controller remained off during this period. During the First Experience Period, after six minutes the exoskeleton controller was turned on for the first time and participants walked for an additional 12 minutes while experiencing the new cost landscape (Fig. 2C). We used this period to determine if participants were *spontaneous initiators* (individuals that adapt toward the optima prior to any perturbation toward higher or lower cost gaits. See *Identifying Spontaneous Initiators* below). We calculated the *first experience preferred step frequency* as the average step frequency during the final 150 seconds of this period. During the Second Experience Period, participants continued to be exposed to the new cost landscape while being held at higher and lower cost gaits (higher and lower step frequencies) by a metronome (Fig. 2D). The metronome tempos were again set to −15% and +10% of initial preferred step frequency to allow participants to experience the extremes of the new cost landscape, while avoiding step frequencies directly to the optima or initial preferred step frequency (approximately −5% and 0% of initial preferred step frequency, respectively). We played each high and low metronome tempo for three minutes, four times each in alternating order, with the first tempo direction randomized. Following each metronome tempo, the metronome turned off for one-minute probes of participants’ self-selected step frequency. We informed participants that at times the metronome would be turned on, during which they should match their steps to the steady-state tempo, and that when the metronome turned off, they no longer had to remain at that tempo. We did not give participants any further directives about how to walk. During the final three minutes of this period, the exoskeleton controller turned off, returning participants to their natural energetic landscape. We calculated the *final preferred step frequency* as the average step frequency during the 150 seconds of the period just prior to the exoskeleton controller turning off. To assess participants’ re-adaptation when returned to the natural cost landscape, we calculated the *re-adaptation preferred step frequency* as the average of the final 150s of the Second Experience Period, after the exoskeleton controller turned off. To determine if participants could articulate an explicit strategy for this energy optimization process, we administered a survey following the final collection period. We asked participants to answer five free form questions (Table 1) in an online platform (Google Forms). We designed these questions to probe their level of awareness and perception of control during optimization.

**Figure 2:**
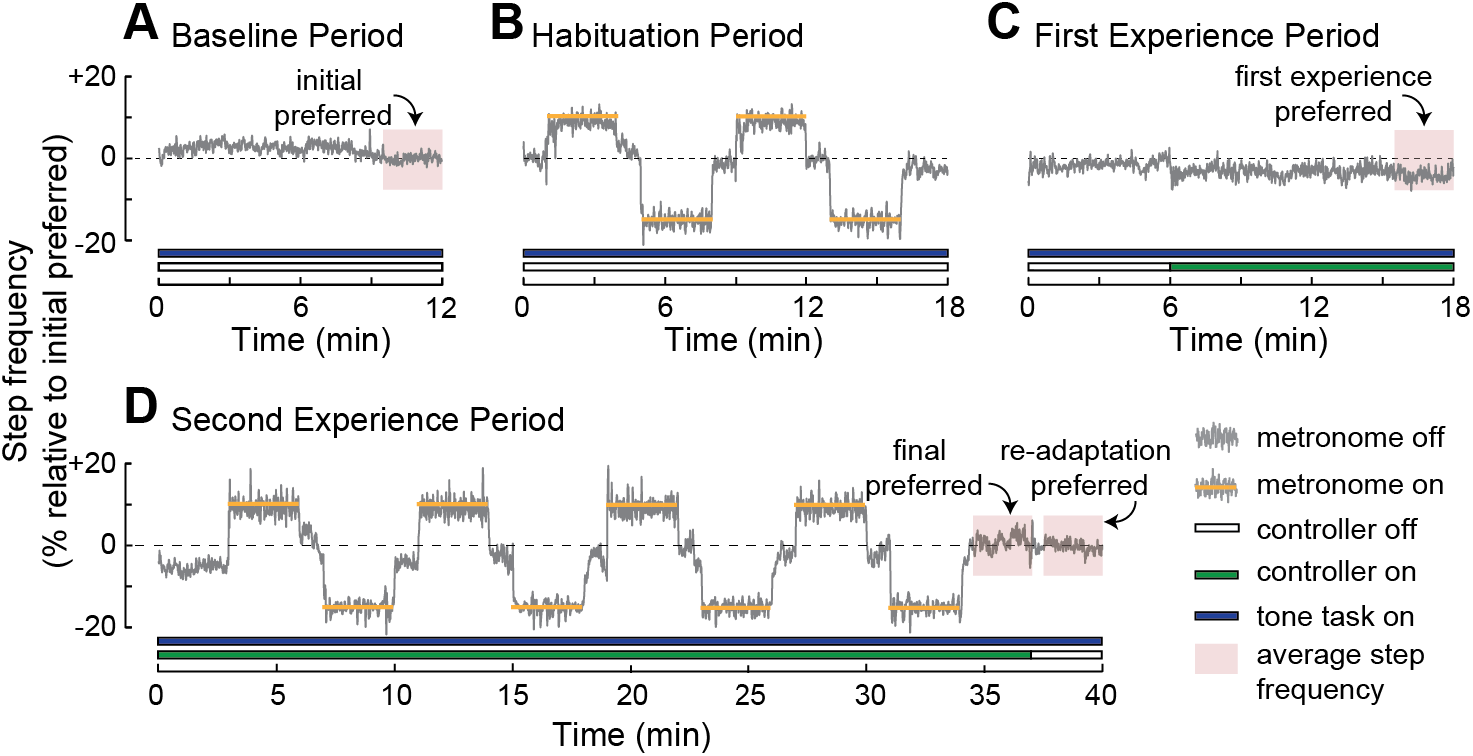
Experimental protocol. Each participant completed four periods: Baseline Period (**A**), Habituation Period (**B**), First Experience Period (**C**), and Second Experience Period (**D**). We provided participants with rest periods of 5–10 minutes between each testing period. Data shown are from one representative participant.

**Table 1:** Survey questionnaire. Participants in the single-task and dual-task experiments answered these five questions in an online form following the final collection period.

| Question # | Keywords* | Question |
| --- | --- | --- |
| 1 | Changed your gait? | When you were walking naturally (no metronome), did you change how you walked? If so, in what way and why? |
| 2 | Made conscious decisions? | When you were walking naturally (no metronome), were you making conscious decisions to change how you walked? If so, how did you make these decisions? And, when did you start making these decisions? |
| 3 | Had control over exo? | Did you feel that you had any control over what the exoskeleton was doing? If so, in what way? |
| 4 | How exo worked? | How was the exoskeleton making walking easier or harder? |
| 5 | Exo walking characteristic? | Did you think any walking characteristic was related to what the exoskeleton was doing? If so, state what characteristic and explain how you thought it related to what the exoskeleton was doing. |

\* Keywords were not provided to participants or response raters but are included here and used in Figure 5 to allow for easier interpretation.

Our dual-task protocol described above does deviate from the previous single-task protocol in a few ways. First, the original single-task experiment used four different metronome tempos (− 15%, −10%, +5% and +10%), while here we used only the two extremes (−15% and +10%). We did so because in our prior experiment we found different effects of probe direction (i.e. +10% vs. −10%), but not magnitudes (i.e. −10% vs. −15%). Here and in other studies subsequent to the single-task experiment (Abram et al. 2019; Selinger et al. 2019; Simha et al. 2019, 2020) we have chosen to simplify our protocol to a single high and low tempo, which are used during both the Habituation Period and the Second Experience Period. Second, in the original single-task experiment, the Second Experience Period was 30 minutes in length, while here it is 40 minutes due to the addition of two extra metronome bouts. This change to a longer Second Experience Period was made to allow us to further investigate the time course of adaptation. However, we have subsequently found that adaptation following metronome holds is largely complete after 20 minutes and so do not expect the protocol change to have a significant effect (Abram et al. 2019). Third, in the original single-task experiment, the exoskeleton controller remained on for six minutes following the last metronome hold, while here in our dual-task experiment it remained on for only three minutes. We made this protocol change to help reduce the total length of the period for participants and because in the original single-task experiment we found adaptation during the final probe to be rapid and complete within tens of seconds (time constant: 10.5 ± 1.8 seconds). When calculating final preferred step frequency in this dual-task experiment, we therefore used a 150-second window of time starting 30 seconds after the final metronome hold, whereas in the original single-task experiment we used a 180-second window of time starting 180 seconds after the final metronome hold. To ensure our primary outcome measure was not affected by this difference, we recalculated the final preferred step frequency from the original single-task experiment data set using the earlier and shorter time window that we use here.

### Experimental outcome measures

To assess performance on our primary energy optimization task, we tested if participants adapted toward the energy optima. To do so, we tested if the average final preferred step frequency decreased from initial preferred step frequency using a one-sample one-tailed *t*-test and a significance level of 0.05. To test if energy optimization was affected by the secondary tone discrimination task, we compared the average final preferred step frequency from our dual-task experiment to that from the previous single-task experiment, calculated over the same time-window. We did so using a two-sample two-tailed *t*-test with a significance level of 0.05. To determine our minimum required participant number we performed an *a priori* power analysis for our primary outcome measure—step frequency adaptation. Based on our two previous studies (Selinger et al. 2015, 2019) we expected complete energy optimization to result in participants decreasing their step frequency by approximately 5% and with an across participant standard deviation of approximately 3.5%, when exposed to the penalize-high controller. To detect an across-participant average difference in step frequency of at least 5%, given an across-participant average standard deviation in step frequency of 3.5% and a single-task participant number of 14, we calculated that we required a minimum of only four dual-task participants to achieve a power of 0.8. Unfortunately, to detect smaller differences in step frequency a prohibitive number of participants would be required. For example, detecting 2.5% or 1% differences would require nearly one-hundred and over ten-thousand participants, respectively. Therefore, in our experiment we chose to test 11 participants, increasing our expected power to over 0.9 when detecting complete versus fully abolished step frequency adaptation. However, it is important to note that we are only able to test if the addition of a secondary explicit task fully disrupts energy optimization.

We also tested if the rate of adaptation was affected by the secondary tone discrimination task. Because most of our participants were non-spontaneous initiators, who required a metronome hold at a low-cost gait before initiating adaptation, we compared the rate of adaptation following the first low holds. Rate of adaptation was calculated by fitting each participant’s step frequency data from one minute prior to the metronome release to the end of the one-minute release period (minutes 5 – 7 of the Second Experience Period) with a single-term exponential curve. To test for differences in adaptation rates between the dual-task and single-task participants, we compared the time constants from the dual-task participants (fit individually and over the same time window) to the reported average time constant from the single-task participants using a one-sample two-tailed *t*-test with a significance level of 0.05.

To assess participants’ re-adaptation when returned to the natural cost landscape, we tested if participants returned to their initial preferred step frequency and the rate at which they converged back to their initial preferred step frequency when the exoskeleton was turned off. To determine whether re-adaptation preferred step frequency values were different from initial preferred step frequency values (0%), we used a one-sample *t*-test with a significance level of 0.05. To assess the rate of re-adaptation, we fit each participant’s step frequency data from the moment the exoskeleton turned off until the end of the Secondary Experience Period (minutes 37 – 40) with a single-term exponential curve. To test for differences between the dual-task and single-task experiments, we compared the time constants from the participants in our dual-task experiment (fit individually and over the same time window) to the reported average time constant from the single-task participants using a one-sample two-tailed *t*-test with a significance level of 0.05.

To assess performance on the secondary tone discrimination task, we calculated response accuracies and reaction times. We calculated these metrics for the same 150-second time windows over which we calculated the initial preferred step frequency during the Baseline Period and the final preferred step frequency during the Second Experience Period. We calculated response accuracy as the percentage of correct button presses during a given time window. We calculated reaction time as the time between presentation of a tone and the onset of a button press, determined by a threshold. To confirm that the secondary tone discrimination task was challenging and required an explicit strategy, we compared participants’ reaction times during the Baseline Period to an average reaction time from a previous experiment where participants completed a simple button press task (Stuss et al. 1989). To do so, we used a one-sample one-tailed *t-*test with a significance level of 0.05. To determine if performance on the secondary tone discrimination task was affected by the primary energy optimization task, we compared reaction times and accuracies during the Baseline Period (when the exoskeleton was off and the energy optima was unchanged) to those during the Second Experience Period (when the exoskeleton was on and the energy optima had shifted to a lower step frequency). We did so using paired-sample *t*-tests with a significance level of 0.05.

To assess participants’ ability to articulate a strategy for the energy optimization process, we compared survey responses from participants in our dual-task experiment to those from the single-task experiment using three independent and blinded raters. All participant responses for a given question, from both the single-task and dual-task experiments, were randomized. Each of the three raters then independently rated all responses for one question (starting with question 1) before moving on to the next. We did not tell raters if the response was from a participant in the single-task experiment or our dual-task experiment. We asked raters to score each response in terms of participant *awareness* and *control* using a 0 to 6 scale. We gave raters the following definitions: ‘*Awareness* refers to the participant’s awareness of the relationship between stride length/step frequency and resistance from the exoskeleton. A rating of 0 means the participant is unaware of the relationship and a rating of 6 means the participant is fully aware. *Control* refers to the participant’s reporting that they changed their stride length/step frequency to control the exoskeleton. A rating of 0 means the participant did not consciously change their gait and a rating of 6 means they did consciously change their gait.’ Raters fully understood our energy optimization paradigm, the exoskeleton control function, and the experimental hypotheses. To compare awareness and control scores between single-task and dual-task participants, we averaged scores across raters to obtain a single score for each participant on each question. We then used a two-sample, one-tailed *t*-test with a significance level of 0.05 to test for differences for each of the five survey questions.

### Identifying spontaneous initiators

Some participants spontaneously initiated optimization, during the First Experience Period, prior to any perturbation to lower or higher cost gaits. To keep our analysis consistent with that from our previous single-task experiment, we tested for the presence of these spontaneous initiators during the First Experience Period and excluded them from all further analyses (Selinger et al. 2019). To identify spontaneous initiators, we used the same two criteria as previously reported. First, the average step frequency at the end of the First Experience Period, first experience preferred step frequency, must be below three standard deviations in steady-state variability, determined from the time window used to calculate initial preferred step frequency. Second, the change in step frequency cannot be an immediate and sustained response to the exoskeleton turning on. To ensure that this was true, first experience preferred step frequency had to be significantly lower than the average step frequency measured from seconds 10 – 40 after the exoskeleton turned on. We tested this using a one-tailed two-sample *t*-test with a significance level of 0.05. We also tested whether there was a difference in the proportion of participants identified as spontaneous initiators between our dual-task experiment and the previous single-task experiment. To do so we used a binomial distribution model with a cumulative distribution function to calculate the probability of identifying at least as many spontaneous initiators as we did in our dual-task experiment, given the prior proportion of spontaneous initiators in the single-task experiment.

## Results

### Identifying spontaneous initiators

We identified three of the eleven participants to be spontaneous initiators. Although this proportion (27%) is higher than that reported in our previous single-task experiment (6/36, 17%), it is not significant (we estimate a 28% chance of finding at least this many spontaneous initiators). On average during the First Experience Period our spontaneous initiators converged to a step frequency lower than their initial preferred step frequency (−3.77% ± 1.31%, p=0.038). In contrast, non-spontaneous initiators remained at a step frequency that was not different from their initial preferred step frequency during this First Experience Period (−1.39% ± 2.25%, p=0.122).

### Tone discrimination task requires explicit attention

Our secondary tone discrimination task was cognitively challenging, demanding attention and explicit processing. We found that even during the Baseline Period, when the exoskeleton controller was off and the energy optima unchanged, participants made response errors. On average response accuracy was 91.4% ± 6.2%, which is better than chance (50%) but not perfect (100%) (Fig. 3A). Moreover, we found that participants’ reaction times were more than three times that typically reported for a simple button press in the absence of a tone discrimination task (0.901s ± 0.073s vs. 0.247 ± 0.014s, p=1.9 × 10^−8^ (Stuss et al. 1989) (Fig. 3B), indicating that the task demanded significant explicit processing.

**Figure 3:**
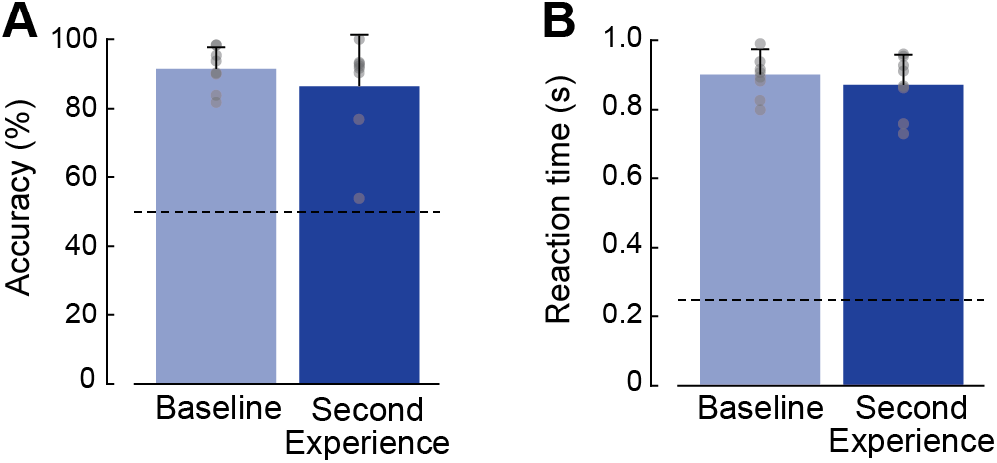
Secondary tone discrimination task performance. **A:** Average accuracy score (%) during the Baseline Period (light blue) and Second Experience Period (dark blue). The dashed horizontal line represents chance (50% accuracy). **B:** Average reaction time (seconds) during the Baseline Period (light blue) and Second Experience Period (dark blue). The dashed horizontal line represents average reaction time (0.247 s) for a simple button press task in the absence of a tone discrimination task (Stuss et al. 1989). Error bars represent one standard deviation. Circles represent individual data from each participant (n=8).

### Tone discrimination task performance was unaffected by the energy optimization process

Participants’ performance on the secondary tone discrimination task did not worsen when the primary energy optimization task was presented. We found no differences in accuracy scores calculated during the Baseline Period, when the exoskeleton was turned off, and the Second Experience Period, when the exoskeleton was turned on and the energy optima changed (91.4% ± 6.2% vs. 86.4% ± 14.7%, respectively; p=0.221; Fig. 3A). The same was true for reaction time scores (0.901s ± 0.073s vs. 0.872s ± 0.086s, respectively; p=0.397; Fig. 3B).

### Tone discrimination task does not disrupt the energy optimization process

Participants optimized their gait to reduce energy expenditure, despite the demands of the secondary tone discrimination task. We found that participants adapted toward the optima, displaying a final preferred step frequency that was lower than their initial preferred step frequency (−3.8% ± 3.5% vs 0%, p=0.010; Fig. 4). Moreover, the magnitude of this adaptation was similar to that for the single-task experiment (−4.02 ± 4.2%; p=0.880). We also found no differences in rate of adaptation between dual- and single-task participants (p=0.149).

**Figure 4:**
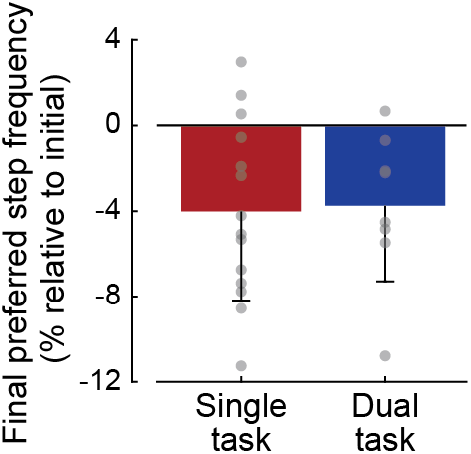
Average final preferred step frequency for participants in the single-task and dual-task experiments (red and blue bars, respectively). Error bars represent one standard deviation. Circles represent individual data from each participant in the single-task and dual-task experiments (n=14 and n=8, respectively).

Furthermore, participants re-adapted to a step frequency similar to their initial preferred step frequency (0%) when the exoskeleton was turned off and they were returned to a natural cost landscape (−1.01% ± 2.3% vs. 0%, p=0.262). The rate of re-adaptation was variable between participants in our dual-task experiment (average time constant: 21.6s ± 25.7s), but we found no differences in rate of re-adaptation between dual- and single-task participants (p=0.262).

### Tone discrimination task disrupts explicit awareness of the energy optimization process

The presence of a secondary explicit task disrupted participants’ awareness of their gait adaptation and perception of control over the exoskeleton during the primary energy optimization task. We found that raters’ average scores of participant awareness in our dual-task experiment were lower than those in the single-task experiment for questions 1 – 3 (0.1 ± 0.2 vs. 2.9 ± 2.0; p=6.6 × 10^−4^, 0.4 ± 1.2 vs. 2.5 ± 2.3; p=0.019, 1.9 ± 2.4 vs. 4.3 ± 2.1; p=0.027; Fig. 5A). For these questions, average scores for our dual-task experiment indicated no to low levels of awareness (scores between 0 and 2), while those for the single-task experiment indicated moderate levels of awareness (scores between 2.5 and 4.5). We found similar differences for raters’ average scores of participant control for questions 1 – 3 (0.1 ± 0.4 vs. 3.8 ± 2.5; p=5.0 × 10^−4^, 0.6 ± 1.6 vs. 3.3 ± 2.6; p=0.014, 1.4 ± 1.6 vs. 4.6 ± 2.1; p=0.002; Fig. 5B). Average scores for our dual-task experiment indicated no to low levels of control (scores between 0 and 1.5), while those for the single-task experiment indicated moderate to high levels of control (scores between 3 and 5). There were no differences in awareness or control scores between dual-task and single-task participants for questions 4 and 5 (Fig. 5A,B).

**Figure 5:**
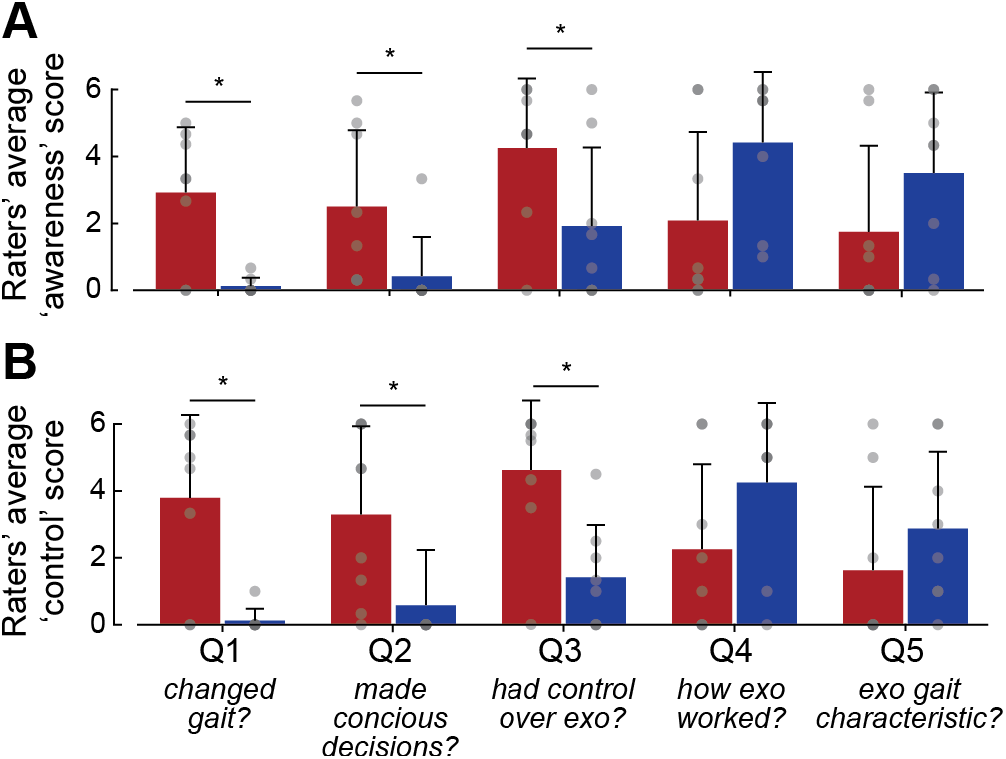
Survey results. **A:** Raters’ average scores of participant awareness. **B:** Raters’ average scores of participants’ perception of control. Red bars represent single-task participants and blue bars represent dual-task participants. Error bars represent one standard deviation. Circles represent individual data from each participant in the single-task and dual-task experiments (n=14 and n=8, respectively).

## Discussion

Here, we used a dual-task paradigm to test whether energy optimization during walking is primarily an implicit or explicit process. We found that adding a secondary, cognitively demanding, explicit task does not disrupt optimization. Participants in our dual-task experiment showed a level of optimization similar to participants in our previous single-task experiment, where attentional resources were not shared with another task. We also found that performance on the secondary tone discrimination task did not worsen when participants were optimizing for energetic cost; accuracy scores and reaction times remained unchanged when the exoskeleton altered the energy optimal gaits. Additionally, the survey responses suggest that dual-task participants were distracted by the secondary task; they were largely unaware of the changes they made to their gait to optimize energy or the control they had over exoskeleton. Interestingly, although single-task participants scored higher for both their awareness of gait change and perception of control, they displayed similar magnitudes and rates of optimization as those in the dual-task. This suggests that even when explicit awareness exists it may not be used during energy optimization. Collectively, our results suggest that energy optimization during walking is primarily an implicit process, requiring minimal conscious attention.

The primary limitation of our experiment is our inability to detect partial changes in the level of optimization between single- and dual-task participants. We found that the magnitude and rate of step frequency adaptations were similar between dual- and single-task experiments—we found no statistical differences. However, variability in individual step frequency measures are high, and although we had the power to detect a full disruption of adaptation (0% vs. 5%), we lacked the statistical power to detect smaller, partial changes. Our results indicate that in our experiment energy optimization is primarily an implicit process, but we are unable to determine if minor explicit contributions existed and were disrupted.

Other limitations of our experiment are inherent to dual-task paradigms. First, it is difficult to know, with certainty, if our participants were sufficiently distracted by the secondary tone discrimination task. Participants’ average accuracy scores (> 85%) were higher than we expected from piloting. If our secondary task was not challenging enough, it is possible that participants had the attentional resources necessary to simultaneously carry out the primary energy optimization task using an explicit process, without displaying performance decrements on either task. However, our survey results suggest this was likely not the case. Dual-task participants were less aware of their gait changes and their ability to affect the exoskeleton behaviour, indicating that they were meaningfully distracted by the secondary task. A second potential interpretation of our findings is that our two tasks draw on distinct cognitive ‘resource pools’. The *central-resource capacity theory* suggests there is a single source of attentional resources for which all simultaneous activities compete, for example walking and having a conversation with a friend (Kahneman 1973; Magill 2011; Schmidt and Lee 2011). Alternatively, *multiple-resource capacity theory* suggests there are several resource pools, each with limited capacity, and each specific to different tasks or processing stages (Magill 2011; Wickens 2010). It is possible that our primary energy optimization task and secondary tone discrimination task draw from two different resource pools, in which case we would not expect performance decrements in either task. Again, however, dual-task participants’ lower survey scores suggest any cognitive awareness of the primary optimization task, whether used to optimize or not, draws from the same pool of resources as our tone discrimination task. Moreover, previous findings, in walking and reaching adaptation paradigms, suggest that tone discrimination tasks can compete for the same resources as these motor tasks (Conradsson et al. 2019; Malone and Bastian 2010; Taylor and Thoroughman 2007). While not possible to conclusively rule out these alternative interpretations, or a minor contribution from an explicit process, our experimental and survey results in combination suggest that energy optimization is primarily an implicit process.

Our distraction task appears to have prevented participants from strategically altering their gait, but may not have fully abolished their explicit understanding of the exoskeleton controller. Our first three survey questions (Q1-3) focused on understanding if participants were strategically changing how they walked (‘Did you change how you walked? Did you make conscious decisions? Did you feel you had control?’). Dual-task participants scored very low on these questions (average scores less than two), and scored significantly lower than those in the single-task experiment (average scores greater than 2.5). Our last two survey questions (Q4-5) focused on understanding if participants understood how the exoskeleton controller worked (‘How did the exoskeleton make walking easier or harder? What gait characteristic affected what the exoskeleton was doing?’). Here, dual-task participants’ average scores were higher than for previous questions (average scores greater than three), although we found no significant difference compared to single-task participants. This suggests that in the presence of the secondary tone discrimination task, participants may still have some explicit understanding of how the exoskeleton controller works, but may not be able to simultaneously develop an explicit gait strategy in response (Bronstein et al. 2009). Single-task participants, who were not distracted and therefore had additional attentional resources, did appear to be able to articulate an explicit gait strategy. That this explicit strategy did not lead to better performance on the energy optimization task—single-task and dual-task participants’ level of gait adaptation and rate of adaptation were similar—further suggests the process of energy optimization itself is primarily implicit.

Our findings are consistent with prior work demonstrating that locomotor adaptations that drive learning are remarkably invariant and unaffected by explicit processes. One approach to understanding the role of conscious control in gait is to disrupt participants’ explicit strategy formation, through a secondary task, and see if this distraction will diminish gait adaptation (as we have done here). An opposite approach is to give participants an explicit strategy, often through direct feedback about the errors they need to reduce, and to see if this awareness will enhance gait adaptation. Malone and Bastian (2010) used both approaches to investigate the role of conscious, or explicit, gait corrections during adaptation to a split-belt treadmill. They found that distraction slowed the rate of adaptation while conscious correction sped it up. However, aftereffects during de-adaptation lasted the longest following distraction, indicating that gait adaptation was more engrained, or better learned, despite the slower adaptation rate. Roemmich et al. (2016) extended this work and demonstrated that explicit information about errors during split-belt walking can lead to rapid and substantial improvements in motor performance without any true improvements in learning. They showed that when explicit feedback is removed, participants revert to a level of gait adaptation consistent with that expected based on rates of adaptation from implicit, in their case proprioceptive, sources. In other words, one can make conscious changes to their gait, based on explicit feedback about errors, but this is not retained and does not improve learning in novel contexts. The finding that voluntary corrections are mechanistically distinct from implicit adaptation and learning is consistent with prior models of gait response to perturbation proposing two processes—one rapid but approximate based on prediction and one slow but accurate driven by optimization (O’Connor and Donelan 2012; Snaterse et al. 2011). That we found no difference in adaptation or de-adaptation between our dual- and single-task experiments implies that in our paradigm, implicit optimization is dominant. It is possible that in other gait adaptation paradigms, such as the split-belt, explicit predictions are more evident because the task is more visually or kinematically clear. The more complex and closed-loop nature of our exoskeleton controller may have prevented rapid explicit prediction. In future, providing participants with explicit feedback about energy costs could offer additional insight into the energy optimization process and serve as an added test of its implicit nature.

That energy optimization is an implicit process has both potential benefits and drawbacks for an adapting human. One clear advantage is that attentional resources can be directed toward other movement objectives. For example, cognitive attention during walking can be directed toward accuracy and navigation demands when encountering obstacles. These explicit demands may act as constraints, while energy optimization proceeds implicitly within these bounds. Another advantage is that when energy optimal solutions are complex and difficult to explicitly predict, the implicit system may still be able to navigate them over sufficient timescales. This may well be the case when people are adapting to injuries that change body mechanics and neural control, or when adapting to assistive devices that apply novel forces and alter limb dynamics, as we did here. While a therapist or prosthetist may be unable to coach an individual to an energy optimal coordination pattern, their nervous system may implicitly learn this over time. However, this ability may be a double-edged sword. Although gait rehabilitation strategies often focus on restoring a desired ‘normal’ or ‘healthy’ gait, our implicit optimization process may be at odds with these kinematic goals if the gaits are no longer energy optimal following injury (Roemmich et al. 2019). A focus on aligning these otherwise competing objectives may lead to more effective and enduring rehabilitation. Another possible disadvantage of implicit optimization is that many have found adaptation that relies solely on an implicit process will be incomplete— even after many trials, residual errors, or an asymptotic offset, persist (Albert et al. 2020; Bond and Taylor 2015). Albert et al. (2020) demonstrated that this offset is a signature of implicit learning and its magnitude relates to one’s sensitivity to past errors. In our experiment, our inability to precisely identify the energy optimal gait makes it difficult to determine if adaptation was incomplete. However, in some of our past work, partial adaptation toward energy optimal gait is clear, further implicating an implicit process during energy optimization (Abram et al. 2019; Simha et al. 2019).

## Acknowledgments

We thank members of the Queen’s Neuromechanics Lab and Simon Fraser Locomotion Lab, as well as Kirsty McDonald for her helpful comments and suggestions.

## Author contributions

Conceptualization: J.C.S., R.L.B., J.M.D.; Methodology: R.L.B., J.C.S.; Formal analysis: R.L.B., M.J.M.; Writing - original draft: M.J.M., J.C.S.; Writing - review & editing: M.J.M., R.L.B., J., J.M.D., J.C.S.; Visualization: M.J.M., J.C.S.; Supervision: J.C.S., J.M.D.; Funding acquisition: J.C.S., J.M.D.

## References

Abram SJ, Selinger JC, Donelan JM. Energy optimization is a major objective in the real-time control of step width in human walking. J Biomech 91: 85–91, 2019.

Albert ST, Jang J, Sheahan H, Teunissen L, Vandevoorde K, Shadmehr R, Albert S, Building T, Hopkins J. An implicit memory of errors limits human sensorimotor adaptation. Pre-print, 2020. doi:10.1101/868406.

Beauchet O, Dubost V, Aminian K, Gonthier R, Kressig RW. Dual-Task-Related Gait Changes in the Elderly: Does the Type of Cognitive Task Matter? [Online]. J Mot Behav 37: 259–264, 2005https://search.proquest.com/docview/216789956?pq-origsite=gscholar&fromopenview=true.

Beauchet O, Kressig RW, Najafi B, Aminian K, Dubost V, Mourey F. Age-related decline of gait control under a dual-task condition. J Am Geriatr Soc 51: 1187–1188, 2003.

Bench CJ, Frith CD, Grasby PM, Friston KJ, Paulesu E, Frackowiakt RSJ, Dolantt RJ. Investigations of the functional anatomy of attention using the Stroop test. Neuropsychologia 31: 907–922, 1993.

Benson BL, Anguera JA, Seidler RD. A spatial explicit strategy reduces error but interferes with sensorimotor adaptation. J Neurophysiol 105: 2843–2851, 2011.

Bond KM, Taylor JA. Flexible explicit but rigid implicit learning in a visuomotor adaptation task. J Neurophysiol 113: 3836–3849, 2015.

Bronstein AM, Bunday KL, Reynolds R. What the “Broken Escalator” Phenomenon Teaches Us about Balance. Ann N Y Acad Sci 1164: 82–88, 2009.

Conradsson D, Hinton DC, Paquette C. The effects of dual-tasking on temporal gait adaptation and de-adaptation to the split-belt treadmill in older adults. Exp Gerontol 125: 110655, 2019.

Ebersbach G, Dimitrijevic MR, Poewe W. Influence of concurrent tasks on gait: A dual-task approach. 1995.

Finley JM, Bastian AJ, Gottschall JS. Learning to be economical: the energy cost of walking tracks motor adaptation [Online]. J Physiol 591: 1081–1095, 2013https://physoc.onlinelibrary.wiley.com/doi/pdf/10.1113/jphysiol.2012.245506 [15 Oct. 2019].

Frensch PA. One concept, multiple meanings: on how to define the concept of implicit learning. In: Handbook of implicit learning, edited by Stadler M, Frensch PA. Thousand Oaks, CA: SAGE Publications Inc., 1998, p. 47–104.

Hagmann-von Arx P, Manicolo O, Lemola S, Grob A. Walking in School-Aged Children in a Dual-Task Paradigm Is Related to Age But Not to Cognition, Motor Behavior, Injuries, or Psychosocial Functioning. Front Psychol 7: 352, 2016.

Kahneman D. Attention and effort. Am J Psychol 88, 1973.

Kahneman D, Egan P. Thiking, Fast and Slow. New York: Farrar, Straus and Giroux, 2011.

Kane MJ, Conway ARA, Miura TK, Colflesh GJH. Working Memory, Attention Control, and the N-Back Task: A Question of Construct Validity. J Exp Psychol Learn Mem Cogn 33: 615–622, 2007.

Kawamura K, Tokuhiro A, Takechi H. Gait analysis of slope walking: a study on step length, stride width, time factors and deviation in the center of pressure. Acta Med Okayama 45: 179–184, 1991.

Lajoie Y, Barbeau H, Hamelin M. Attentional requirements of walking in spinal cord injured patients compared to normal subjects [Online]. Spinal Cord 37: 245–250, 1999http://www.stockton-press.co.uk/sc [22 Apr. 2020].

Lajoie Y, Teasdale N, Bard C, Fienry M. Attentional demands for static and dynamic equilibrium. Springer-Verlag, 1993.

Li K, Lindenberger U, Marsiske M, Baltes PB. Memorizing while walking: Increase in dual-task costs from young adulthood to old age. Psychol Aging 15: 417–436, 2000.

Magill RA. Attention as a limited capacity resource. In: Motor Learning and Control: Concepts and Applications. New York: McGraw-Hill Companies, 2011, p. 194–220.

Malone LA, Bastian AJ. Thinking About Walking: Effects of Conscious Correction Versus Distraction on Locomotor Adaptation. J Neurophysiol 103: 1954–1962, 2010.

Masters RSW, Poolton JM, Maxwell JP, Raab M. Implicit Motor Learning and Complex Decision Making in Time-Constrained Environments. J Mot Behav 40: 71–79, 2008.

Mazaheri M, Roerdink M, Bood RJ, Duysens J, Beek PJ, Peper CLE. Attentional costs of visually guided walking: Effects of age, executive function and stepping-task demands. Gait Posture 40: 182–186, 2014.

Mazzoni P, Krakauer JW. An implicit plan overrides an explicit strategy during visuomotor adaptation. J Neurosci 26: 3642–3645, 2006.

Montero-Odasso M, Muir SW, Speechley M. Dual-task complexity affects gait in people with mild cognitive impairment: The interplay between gait variability, dual tasking, and risk of falls. Arch Phys Med Rehabil 93: 293–299, 2012.

O’Connor SM, Donelan JM. Fast visual prediction and slow optimization of preferred walking speed. J Neurophysiol 107: 2549–2559, 2012.

Paul SS, Ada L, Canning CG. Automaticity of walking-implications for physiotherapy practice. Phys Ther Rev 10: 25–23, 2005.

Peper CLE, Oorthuizen JK, Roerdink M. Attentional demands of cued walking in healthy young and elderly adults. Gait Posture 36: 378–382, 2012.

Regnaux JP, David D, Daniel O, Ben D, Combeaud M, Bussel B. Evidence for Cognitive Processes Involved in Control of Steady State of Walking Evidence for Cognitive Processes Involved in the Control of Steady State of Walking in Healthy Subjects and after Cerebral Damage. Neurorehabil Neural Repair 19: 125:132, 2005.

Roemmich RT, Leech KA, Gonzalez AJ, Bastian AJ. Trading Symmetry for Energy Cost During Walking in Healthy Adults and Persons Poststroke. Neurorehabil Neural Repair 33: 602–613, 2019.

Roemmich RT, Long AW, Bastian AJ. Seeing the Errors You Feel Enhances Locomotor Performance but Not Learning. Curr Biol 26: 2707–2716, 2016.

Schmidt R, Lee TD. Motor Control and Learning. 5th ed. Windsor, ON: Human Kinetics, 2011.

Selinger JC, O’Connor SM, Wong JD, Donelan JM. Humans Can Continuously Optimize Energetic Cost during Walking. Curr Biol, 2015. doi:10.1016/j.cub.2015.08.016.

Selinger JC, Wong JD, Simha SN, Donelan JM. How humans initiate energy optimization and converge on their optimal gaits. J Exp Biol 222: jeb198234, 2019.

Shadmehr R, Mussa-Ivaldi FA. Adaptive representation of dynamics during learning of a motor task. J Neurosci 14: 3208–3224, 1994.

Shadmehr R, Smith MA, Krakauer JW. Error Correction, Sensory Prediction, and Adaptation in Motor Control. Annu Rev Neurosci 33: 89–108, 2010.

Simha SN, Wong JD, Selinger JC, Abram SJ, Maxwell J. Increasing the gradient of energetic cost does not initiate adaptation in human walking. Pre-print, 2020. doi:10.1101/2020.05.20.107250.

Simha SN, Wong JD, Selinger JC, Donelan JM. A Mechatronic System for Studying Energy Optimization During Walking. IEEE Trans Neural Syst Rehabil Eng 27: 1416–1425, 2019.

Snaterse M, Ton R, Kuo AD, Maxwell Donelan J. Distinct fast and slow processes contribute to the selection of preferred step frequency during human walking. J Appl Physiol 110: 1682–1690, 2011.

Sparrow WA, Bradshaw EJ, Lamoureux E, Tirosh O. Ageing effects on the attention demands of walking. Hum Mov Sci 21: 961–972, 2002.

Stroop Y. The basis of Ligon’s theory. Am J Psychol 47: 499–504, 1935.

Stuss DT, Stethem LL, Hugenholtz H, Picton T, Pivik J, Richard MT. Reaction time after head injury: Fatigue, divided and focused attention, and consistency of performance. J Neurol Neurosurg Psychiatry 52: 742–748, 1989.

Sun J, Walters M, Svensson N, Lloyd D. The influence of surface slope on human gait characteristics: a study of urban pedestrians walking on an inclined surface. Ergonomics 39: 677–692, 1996.

Taylor JA, Krakauer JW, Ivry RB. Explicit and implicit contributions to learning in a sensorimotor adaptation task. J Neurosci 34: 3023–3032, 2014.

Taylor JA, Thoroughman KA. Divided attention impairs human motor adaptation but not feedback control. J Neurophysiol 98: 317–326, 2007.

Theill N, Martin M, Schumacher V, Bridenbaugh SA, Kressig RW. Simultaneously measuring gait and cognitive performance in cognitively healthy and cognitively impaired older adults: The basel motor-cognition dual-task paradigm. J Am Geriatr Soc 59: 1012–1018, 2011.

Weerdesteyn V, Schillings AM, Duysens J, Van Galen GP. Distraction Affects the Performance of Obstacle Avoidance During Walking. J Mot Behav 35: 53–63, 2003.

Wickens CD. Multiple resources and performance prediction. Theor Issues Ergon Sci 3: 159–177, 2010.

Willingham DB. A neurophysiological theory of motor skill learning. Psychol Rev 105: 558–584, 1998.

Woollacott M, Shumway-Cook A. Attention and the control of posture and gait: A review of an emerging area of research. Gait Posture 16: 1–14, 2002.

